# *Streptococcus mutans LiaS* gene regulates its cross-kingdom interactions with *Candida albicans* to promote oral candidiasis

**DOI:** 10.1101/2025.02.10.637462

**Authors:** Yanfan Cao, Shan Huang, Jingyun Du, Xiuqing Wang, Guowu Gan, Shuai Chen, Xiaojing Huang

## Abstract

*Candida albicans* and *Streptococcus mutans* are commonly associated with oral candidiasis and dental caries, respectively. *C. albicans* is frequently detected in the dental plaque of early childhood caries. The synergistic interaction between *C. albicans* and *S. mutans* in dental caries suggests that fungi enhance the production of acidic metabolites and contributes to the development of caries. However, the cross-kingdom interaction of *S. mutans* in oral candidiasis and the pathways regulating hyphal formation remain unclear. The *liaS* gene of *S. mutans* 593 plays a crucial role in maintaining membrane homeostasis. In comparison to the *S. mutans* 593 and liaS^-^ complementary strains, the liaS^-^ strain significantly diminished the proliferation, biofilm formation, extracellular polysaccharide production, and *C. albicans* hyphal transformation of co-cultured microorganisms. The deletion of *liaS* gene in the dual-species downregulated the expression of yeast-hyphal transformation related genes *hwp1, ece1, als1*, and *als3* in *C. albicans*, while also impacted the expression of *phr1* and *phr2*. Additionally, liaS^-^ inhibited glycolysis and pyruvate kinase activity in *S. mutans*, leading to a decreased in water-soluble and water-insoluble glucans production in biofilm. The significant downregulation of glucosyltransferase transcription relates genes expression in liaS^-^ resulted in inhibition of the cross-kingdom interaction in the dual-species. In the murine oropharyngeal candidiasis model, dual-species infection with liaS^-^ results in a reduction of fungal and hyphal invasion compared to the combinations of *C. albicans* and *S. mutans* 593. In conclusion, *S. mutans*/*liaS* play a crucial role in the development of oral candidiasis suggests that it could be a targeted strategy for managing this condition.

**Importance:** *Candidiasis* involves the transition of *C. albicans* into hyphae and the fungi invasion of the oropharyngeal mucosa. The transformation of hyphal is influenced by a variety of internal and external environmental factors. Our research findings suggest that the deletion of the *liaS* gene leads to a reduction in *gtfs* gene expression, impacting glycolysis efficiency and effectively decreasing EPS production in the dual-species biofilm. Also, downregulation of the expression levels of yeast-hyphal transformation related genes to inhibiting the virulence function of *C. albicans*. Our study demonstrates that *S. mutans* 593/*liaS* promotes epithelial invasion of *C. albicans* in mouse oropharyngeal candidiasis, offering a potential therapeutic approach for addressing cross-kingdom infections.

## Introduction

The rising incidence of fungal infections, especially their effects on human health and quality of life, has emerged as a major concern and poses a significant challenge for clinical management (1). In recent decades, the rapid increase in patients with systemic diseases such as cancer and AIDS, advancements in treatment modalities, and the emergence of drug resistance have all contributed to a rapid surge in cases of fungal infections (2). Candidiasis is a prevalent oral fungal infection that can have a substantial impact on localized lesions or underlying systemic diseases (3, 4). Recurrent infections in individuals with oral candidiasis may suggest compromised host immune defenses, resulting in mucosal invasion and the formation of white plaques (5, 6). In certain instances, recurrent candidiasis may be linked to a heightened susceptibility to developing oral cancer in genetically predisposed individuals (7).

*Candida albicans* (*C. albicans*), the primary causative agent of oral candidiasis, demonstrates a colonization rate exceeding 50% in the human oral cavity, which increases to over 80% in individuals afflicted with the disease (8, 9). This fungus exhibits three distinct morphological forms: yeast, pseudo-hyphae, and hyphae, with hyphal transformation playing a critical role in its adherence and invasive capabilities. During hyphal development, *C. albicans* releases a variety of enzymes that compromise oral epithelial integrity and disrupt cellular cross-linking while simultaneously triggering the host’s immune responses (10). In addition, it synthesizes adhesion-related proteins, particularly from the agglutinin-like sequence (*als*) gene family, which facilitate adherence to epithelial cells, symbiotic microorganisms, and various pattern recognition receptors (11). The transition from yeast to hyphae is intricately regulated by a complex interplay of internal and external factors. Internal regulation involves the expression of hyphal wall protein (*hwp*) relate gene and enhanced candida toxin relate gene (*ece1*), to regulate the synthesis of metabolites required for germ-tube growth. The environmental factors such as acidity, temperature, nutrition, and other microorganisms also play a role in hyphal transformation and biofilm formation. Targeted inhibition of hyphal transformation presents a promising therapeutic strategy for controlling *C. albicans* infections and mitigating drug resistance (12, 13).

*Streptococcus mutans* (*S. mutans*) and *C. albicans* have been shown to enhance biofilm formation, rapid binding, and spreading characteristics when cultured in vitro, resulting in a 3D structured cross-kingdom combination. This phenomenon has been confirmed through the identification of a high proportion of *C. albicans* in early childhood caries mediated by *S. mutans* (14, 15). The secretion of Als3 protein on the surface of *C. albicans* promotes bacterial adhesion and copolymerization, leading to increased production of superoxide dismutase 1 (*sod1*) for enhanced bacterial oxidative stress resistance. Furthermore, the *phr1* and *phr2* genes adapt to environmental pH regulation, facilitating acid production and extracellular matrix production to augment the cariogenic potential of *S. mutans* (16-18). Glycolysis is sensitive to acidic environments, leading to synergistic interactions. It catalyzes the breakdown of sucrose into glucose and fructose, thereby promoting the utilization and growth of sugar sources by *C. albicans* (19). Additionally, glucosyltransferases play a role in promoting binding and extracellular polysaccharide (EPS) production, thereby enhancing the biofilm structure. This cross-kingdom combination strengthens the biofilm structure and promotes cariogenic activity and hyphal invasion (20, 21).

The two-component signal (TCS) transduction system regulates the growth, response to environmental changes, and microbial competition of *S. mutans* through the acid tolerance response. TCS is composed of a transmembrane sensor histidine kinase and a response regulator (22). *LiaSR* is among the 15 TCSs and is involved in regulating bacterial responses to environmental stress, cell membrane damage, and drug resistance (23). Our group previously discovered that *liaS* gene modulates the acid tolerance response of *S. mutans* and carcinogenicity (24). Clinical high caries patients (DMFT ≥ 6) colonized with serotype C *Streptococcus mutans* 593 (*S. mutans* 593) exhibit heightened sensitivity to the regulation of *liaS* gene. However, the role of *liaS* in the cross-kingdom interaction with *C. albicans* and its contribution to the development and progression of oral candidiasis remains poorly understood. This study aimed to elucidate the function of *S. mutans*/*liaS* in modulating its cross-kingdom interaction with *C. albicans* within a dual-species biofilm as well as in a murine oropharyngeal candidiasis model, while also investigating the mechanism by which *liaS* gene regulates hyphal transformation.

## Results

### Effect of *liaS* deletion on chain-length in *S. mutans*

The plasmid *liaS*/pIB169 was utilized for integrating the *liaS* gene of liaS^-^ to generate the complementary strain designated as *S. mutans* 593-liaS^-^-comp. The chain-length in the *liaS* deletion strain was observed to be shorter compared to that of the control strain *S. mutans* 593. Upon supplementation of *liaS*, the chain-length was restored (Supplementary Fig. 2A). qRT-PCR results demonstrated expression of the liaS gene in several screened strains (Supplementary Fig. 2B), and pulse-field gel electrophoresis analysis revealed expression of chromosomal DNA in complementary strains (Supplementary Fig. 2C).

### *LiaS* regulates its cross-kingdom interaction with *C. albicans*

Through the crystal violet (CV) staining, it was discovered that *S. mutans* and *C. albicans* synergistically promoted biofilm formation. Compared with *S. mutans* 593 or liaS^-^-comp, *liaS* plays a crucial role in the biofilm formation of single and dual-species (Fig. 1A). This indicated that the *liaS* gene is involved in regulating the formation of biofilms in *S. mutans* and its combination with *C. albicans*. Compared to the cultivation of *C. albicans* alone, the fungal biomass in the dual-species exhibited only a slight increased (Fig. 1B). Moreover, the biomass of *S. mutans* showed a significant increased that suggesting an enhanced colonization ability of *S. mutans* in cross-kingdom combination biofilms (Fig. 1B). However, when the *liaS* gene is deleted, the colonization ability is markedly reduced, which is consistent with its biofilm formation.

**Figure 1.**
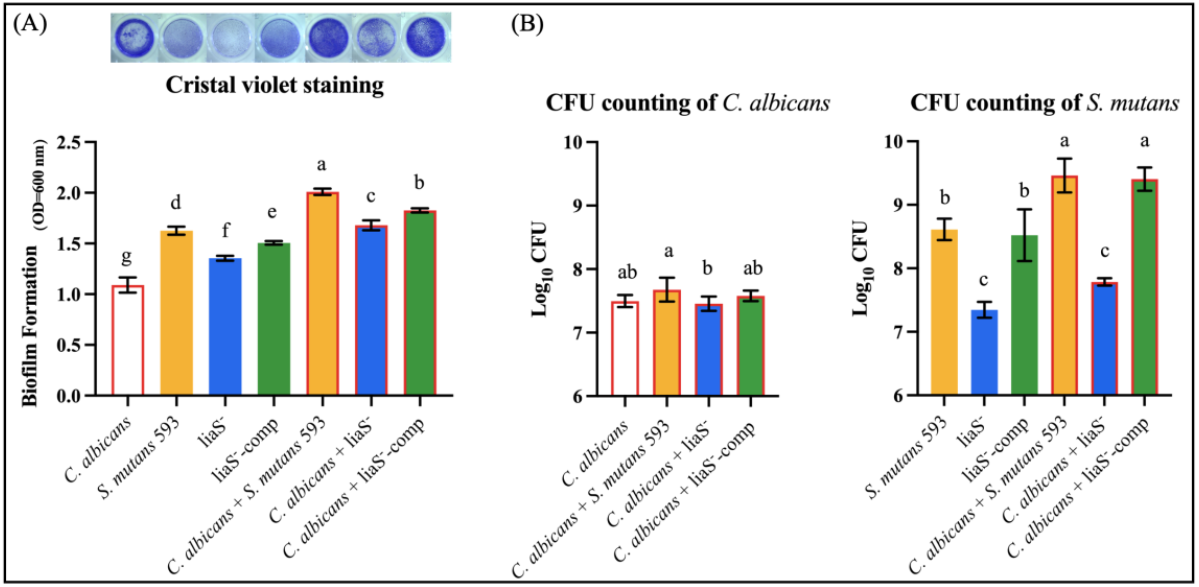
Single and dual-species of *S. mutans* and *C. albicans* biofilm biomass determined by crystal violet assay and CFU count. (A) Quantitative analysis of biofilm biomass by crystal violet assay with inserted panel of representative single and dual-species biofilm stained with crystal violet. The result showed obvious inhibit effect of *liaS* gene on single and dual-species biofilms; (B) CFU count of *S. mutans* and *C. albicans* in biofilms after 24 h. Data were presented in mean ± standard deviation and values with dissimilar letters were significantly different from each other; *P* < *0*.*05*.

### *LiaS* effect biofilm structure and fungal morphology

Scanning electron microscopy (SEM) was utilized to observe the biofilm to further investigate the impact of *S. mutans*/*liaS* on its biofilm structure and fungal morphology with *C. albicans*. The biofilm of *C. albicans* exhibited uniform distribution without evident EPS, with several germ tubes extending to connect with additional cells (Fig. 2). Single-species *S. mutans*, including *S. mutans* 593, liaS^-^, and liaS^-^-comp, formed a biofilm consisting of densely aggregated bacterial colonies and EPS. In comparison to *S. mutans* 593 and liaS^-^-comp, liaS^-^ displayed characteristics of lower self-assembled aggregation, smaller and more dispersed colonies, as well as reduced EPS production within the biofilm. The cross-kingdom combination of *C. albicans* and *S. mutans* resulted in a dense and combined complex structure. When combined with *S. mutans* 593 or liaS^-^-comp, the two microorganisms adhered to each other forming a dense reticulate structure through hyphae; however, the proportion of *S. mutans* adhered to *C. albicans* in the combined biofilm with liaS^-^ is relatively lower while fungi mainly existed in yeast form.

**Figure 2.**
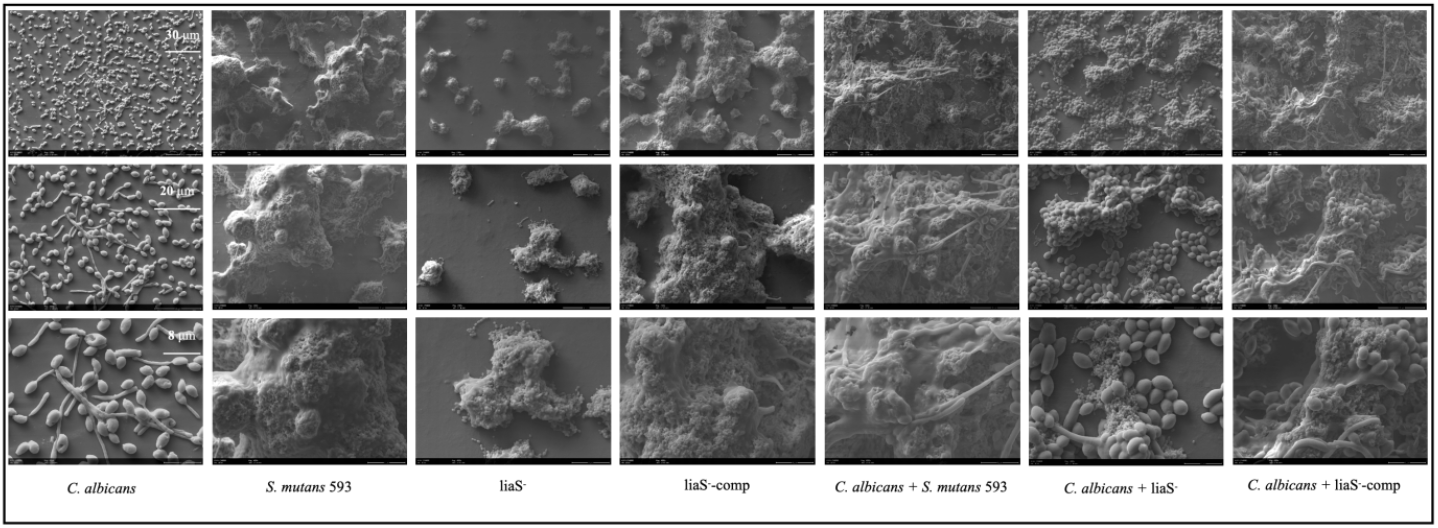
The structures and fungal morphology of cross-kingdom biofilms were influenced by *liaS* gene. Scanning electron microscopy images of 24 h biofilms revealed that a greater number of yeasts were present in the *C. albicans* + liaS^-^ combination, while fewer yeasts but additional hyphae of fungal cells were observed in *C. albicans* + *S. mutans* 593 or liaS^-^-comp. The images were captured at magnifications of 1,000×, 2,000×, and 4,000×. Representative images from at least five randomly chosen samples are presented, with scale bars indicating measurements of 30 μm, 20 μm, and 8 μm.

### *LiaS* promotes glycolysis and EPS production

The 3-(4,5-dimethylthiazol-2-yl)-2,5-diphenyl tetrazolium bromide (MTT) assay of biofilm revealed that the deletion of *liaS* had a significant impact on reducing metabolism compared to *S. mutans* 593 or liaS^-^-comp (Fig. 3A). By monitoring the decrease in glycolytic pH of single-species and dual-species, it revealed that the *liaS* gene is closely associated with the acid production function. *S. mutans* serves as the main functional role in dual-species glycolysis, and *C. albicans* can enhance the glycolysis efficiency to it. In the *liaS* gene deleted strain, the glycolytic pH value decreased at a lower rate (Fig. 3B). pyruvate kinase (PK) is one of the most crucial enzymes involved in EPS and acid production, and its activity also indicates the cariogenic potential of *S. mutans*. Further tests demonstrated that liaS^-^ inhibited PK activity in the biofilm (Fig. 3C).

**Figure 3.**
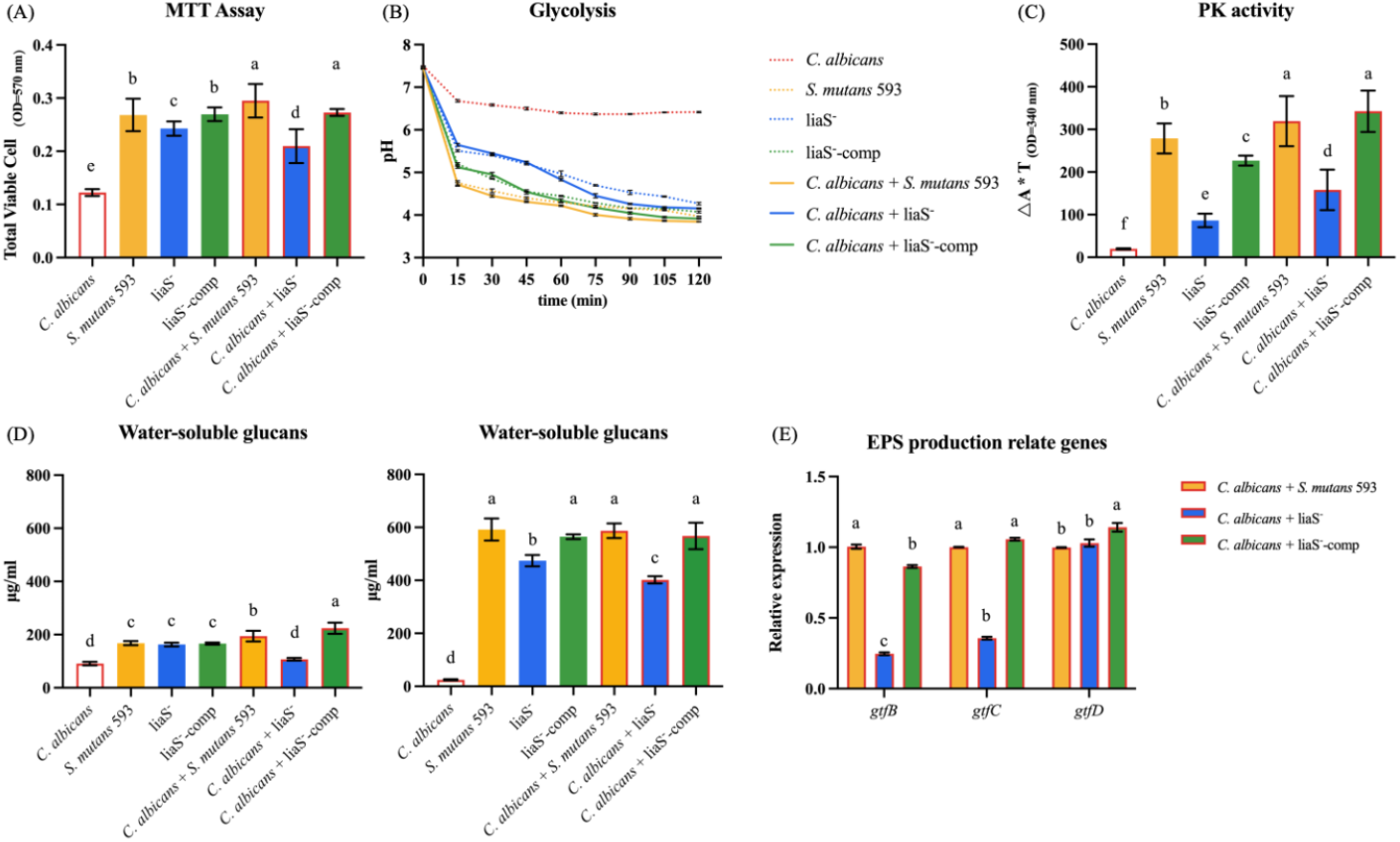
Metabolic activity and EPS production of *S. mutans* and *C. albicans* biofilms. This included: (A) MTT assay of single-species and dual-species biofilm; (B)/(C) The pH drop result reflects the efficiency of glycolysis, and the activity of PK is the main regulatory pathway of glycolysis; (D) Analysis of water-soluble and water-insoluble glucans in single-species and dual-species biofilms, as EPS is primarily composed of these polysaccharides; (C) Investigation into the relationship between EPS production and gene expression of *S. mutans* in cross-kingdom biofilms. The data presented are shown as mean ± standard deviation. Data were presented in mean ± standard deviation and values with dissimilar letters were significantly different from each other; *P* < *0*.*05*.

The production of water-soluble and water-insoluble glucans in biofilms has been shown to play a crucial role in promoting the hyphal transformation of *C. albicans* by *S. mutans* in cocultured biofilms through EPS production. Mutation of liaS resulted in a substantial decrease in the production of both EPS in biofilms (Fig. 2D). This effect appeared to be more pronounced in the dual-species biofilm, and the combination of *C. albicans* and liaS^-^-comp restored the expression of these functions, indicating that liaS has an impact on the cross-kingdom interaction between *S. mutans* and *C. albicans*. We monitored genes involved in EPS production, including *gtfB, gtfC*, and *gtfD* (Fig. 2E). In summary, within the cocultured biofilm of liaS^-^, there was a significant downregulation in the expression of *gtfB* and *gtfC*, while the expression of *gtfD* remained unchanged (*P > 0*.*05*).

### Cross-kingdom effect of *lias* gene on hyphal formation

Observed the yeast to hyphae transformation in germ tubes (GT) formation fungi under an inverted optical microscope at 40× for cell counting analysis. The proportion of hyphae was found to be higher in the combination of *C. albicans* + *S. mutans* 593 and *C. albicans* + liaS^-^-comp compared to the combination of *C. albicans* and *C. albicans* + liaS^-^ (Fig. 4A). The GT formation counts results indicated a decrease by 65.3% for *C. albicans* single-species and decrease by 52.6% for *C. albicans* + liaS^-^, compared to *C. albicans* + *S. mutans* 593 and *C. albicans* + liaS^-^-comp (*P* < *0*.*05*; Fig. 4B). Additionally, it was observed that the complementary strain did not have a significant effect on GT formation. These findings further support the role of *S. mutans*/*liaS* in regulating the hyphal transformation of cross-kingdom with *C. albicans*.

**Figure 4.**
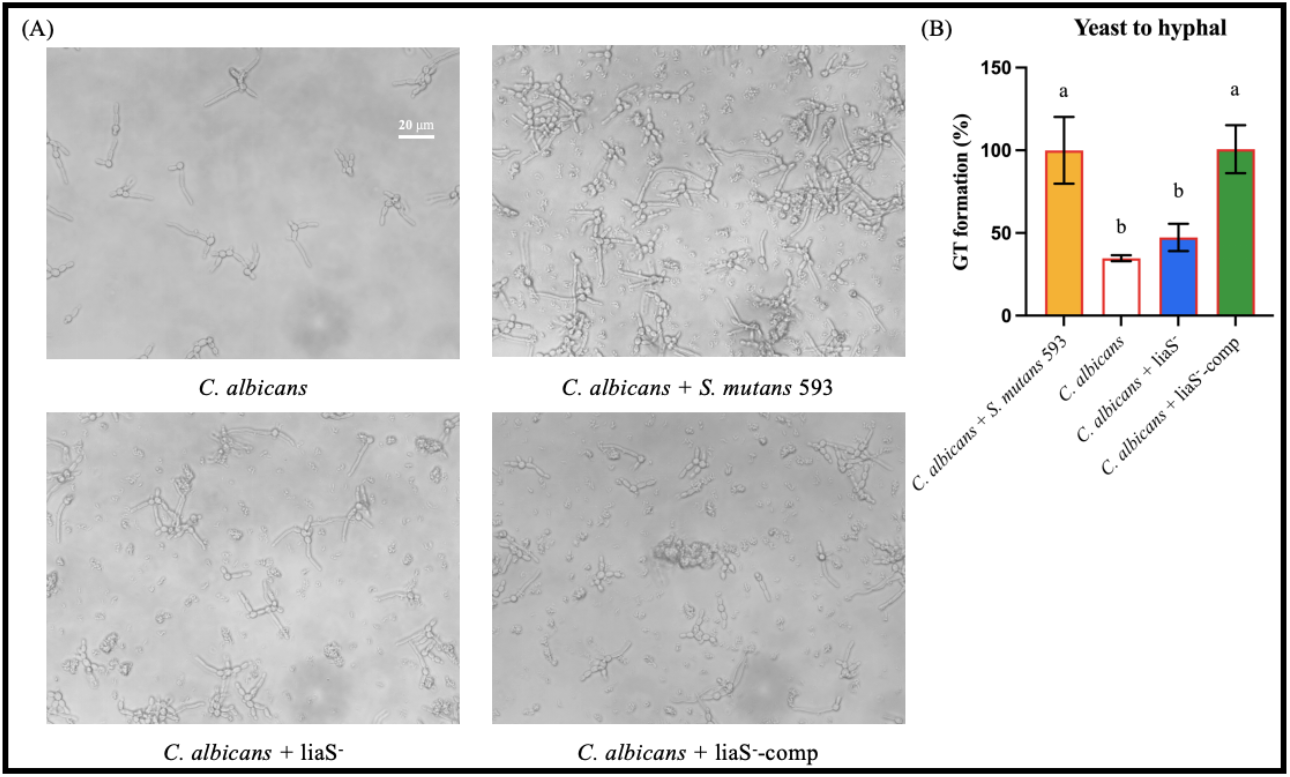
The single-species and *liaS* gene deleted mutation of dual-species reduced the yeast to hyphal transformation by 65.3% and 52.6%, respectively, compared to the combination of *C. albicans* coculture with *S. mutans* 593, and set as the control group. Data were presented as mean ± standard deviation, with dissimilar letters were significantly different from each other; *P* < *0*.*05*.

The impact of *liaS* on the *C. albicans* cell membrane in cross-kingdom biofilms was assessed using an ergosterol biosynthesis assay. Both single and dual-species biofilms exhibited characteristic peaks of ergosterol in the spectrum (Fig. 5A). However, the level of ergosterol biosynthesis reflects the growth of *C. albicans*. The ergosterol content in fungi decreased to 21.2% and 5.3%, respectively, compared to *C. albicans* + *S. mutans* 593 (Fig 5B).

**Figure 5.**
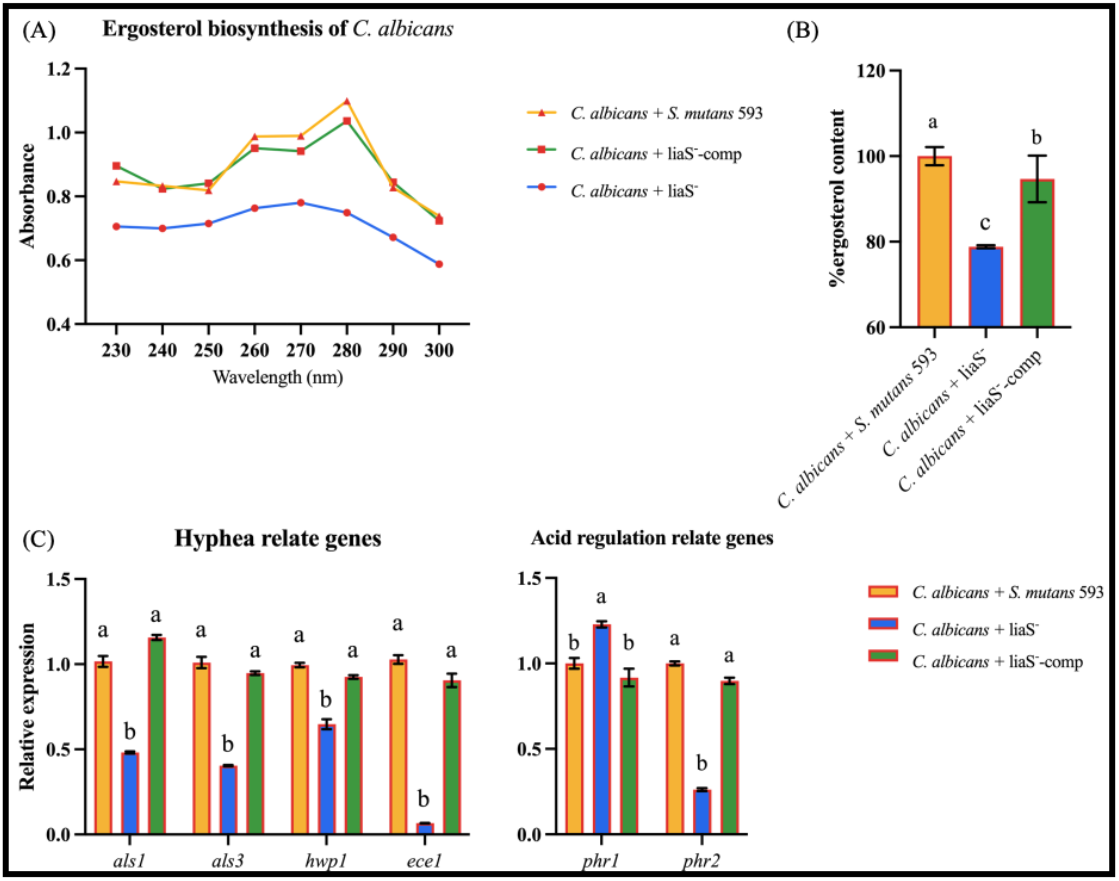
The impact of *liaS* gene on ergosterol biosynthesis was assessed by measuring the spectral profiles of ergosterol extracted from *C. albicans* cells using n-heptane. (A) The results showed a significant reduction in ergosterol content in both single-species and *C. albicans* + liaS^-^, as demonstrated in the plot. (B) Formula calculations further confirmed the decrease in ergosterol content in *C. albicans* + liaS^-^ and *C. albicans* + liaS^-^-comp compared to the control group, *C. albicans* + *S. mutans* 593. Virulence associated with the gene expression of *C. albicans* in cross-kingdom biofilms. Adhesion and hyphal transformation are related to the gene expression of *C. albicans* in cross-kingdom biofilms. Additionally, there is a correlation between the genes of *C. albicans* and their response to pH changes in cross-kingdom biofilms, where *phr1* expression increases at pH > 5.5 and *phr2* expression increases at pH < 5.0. The data were presented as mean ± standard deviation, with dissimilar letters were significantly different from each other; *P* < *0*.*05*.

### Hyphae transformation relates genes expression

The formation of hyphae in *C. albicans* is essential for its virulence. We aimed to further investigate the impact of *S. mutants*/*liaS* on the pathogenicity of *C. albicans* by examining the expression of *als1, als3, hwp1, ece1, phr1* and *phr2* genes involved in hyphal development and adaptation to acidic environments. The co-culture of *C. albicans* and *S. mutans* 593 or liaS^-^-comp significantly upregulated genes related to adhesion and hyphal transformation in *C. albicans* biofilms, while *C. albicans* + liaS^-^ had a significant inhibitory effect (*P* < *0*.*05*; Fig. 5C). This aligns with the observed inhibition of *gtfB* and *gtfC* expression as well as reduced glucan production by *S. mutans*/*liaS*, indicating a cross-kingdom interaction affecting *C. albicans*’ hyphal transformation through glucosyltransferase (Gtfs) and EPS biosynthesis regulation pathway (Fig. 3E). The EPS content showed an inverse correlation with microenvironment pH levels, suggesting that pH changes can influence hyphal transformation process. The cell wall glucan remodeling enzyme Phr has been found to impact the ability of *C. albicans* to adapt to environmental pH. Specifically, *phr1* expression increases in environments with a pH > 5.5, while *phr2* expression increases in environments with a pH < 5.0. These changes in *phr* gene expression have been shown to affect hyphal transformation. Interestingly, qRT-PCR results revealed a significant increase in the expression of *phr1* and a significant decrease in the expression of *phr2* in *C. albicans* + liaS^-^ (*P* < *0*.*05*; Fig. 5C). The pathway through which *liaS* promotes hyphal transformation may be related to glycolysis regulation.

### *LiaS* gene promoted pathogenicity of *C. albicans* in murine oropharyngeal candidiasis

The murine oropharyngeal candidiasis model was utilized to demonstrate the promotion of mucosal invasion of *C. albicans* by *S. mutans*/*liaS* (Fig. 6A). Mice infected with *C. albicans* exhibited significant weight loss, with reductions of 7.8%, 15.3%, and 8.8% compared to their initial weight, respectively (Fig. 6B). *S. mutans*/*liaS* has increased fungal burdens in tongue, stomach, and kidney of infected mice (*P* < *0*.*05*; Fig. 6C). Furthermore, observation of the tongues of mice infected with fungi revealed white patches and several papillary masses in the tongue with *C. albicans* + *S. mutans* 593, while no significant pathological changes in the epithelial regions were observed in *C. albicans* and *C. albicans* + liaS^-^ (Fig. 8D).

**Figure 6.**
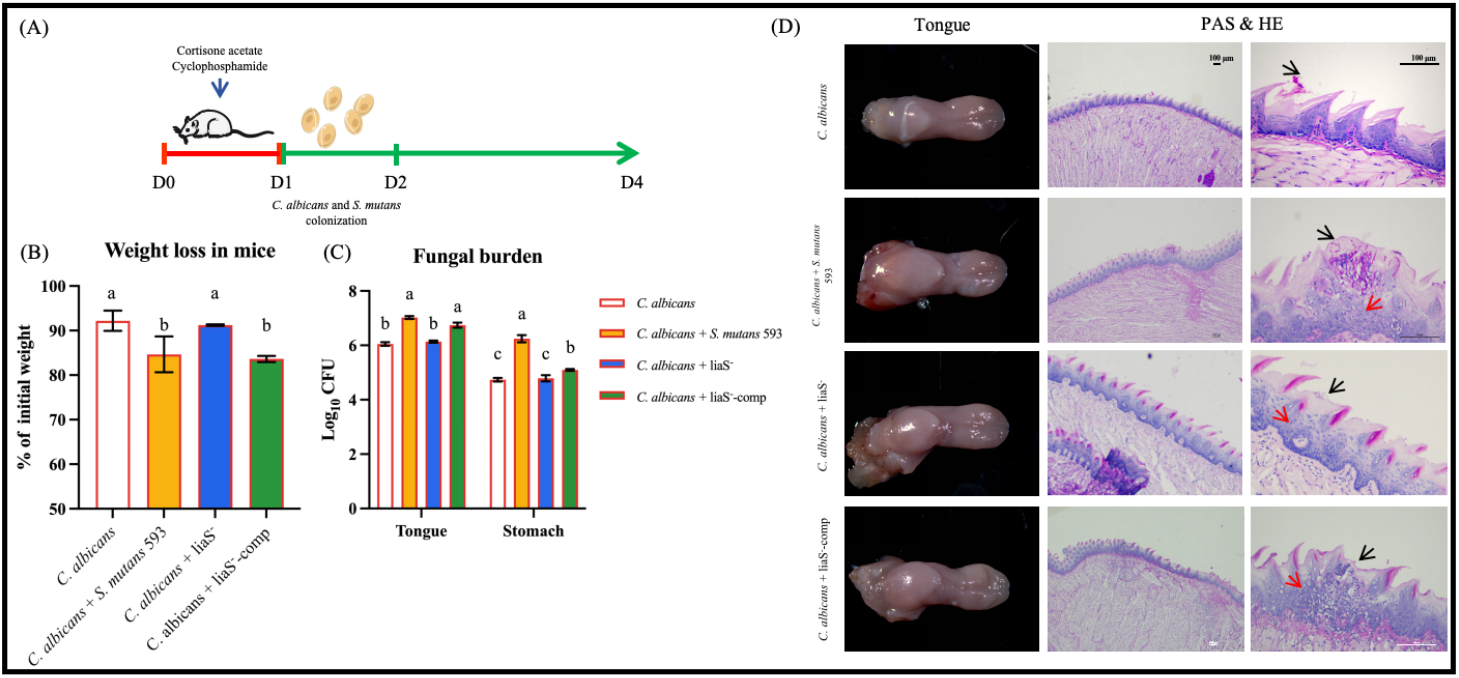
(A) Schematic representation of murine oropharyngeal candidiasis model. (B) The ratio of weight to initial in mice after 3-day infection with *C. albicans*. (C) Fungal burden following 3-day infection with *C. albicans. C. albicans* cells were isolated from the tongues, and stomach of mice. Data were presented as mean ± standard deviation, with dissimilar letters were significantly different from each other; *P* < *0*.*05*. (D) Images showing, HE and PAS-stained tongues, as well as 40× and 200× magnification views of the tongues. Inflammatory cells are indicated by red arrowheads, while invading hyphae are indicated by black arrowheads. HE: haematoxylin/eosin; PAS: Periodic acid–Schiff

Subsequently, the histological and pathological analysis of *C. albicans*-infected tongues revealed varying degrees of invasion and inflammation in immunodeficient mice. Fungal colonization resulted in thickening of the corneal layer, infiltration of neutrophils and macrophages, as well as focal necrosis of the mucosal epithelium. However, *C. albicans* and liaS^-^ mice showed lower fungal invasion and minimal inflammatory cell infiltration on the tongue tissue surface. The Periodic acid-Schiff (PAS) staining confirmed that *C. albicans*’ invasion, with hyphal colonization leading to germ invasion and necrotic cell aggregation in combination with *C. albicans* + *S. mutans* 593. In contrast, *C. albicans* and *C. albicans* + liaS^-^ exhibited significantly reduced invasion degree, consistent with the regulatory ability of the *liaS* gene on hyphal transformation in vitro.

## Discussion

*S. mutans* and *C. albicans* are known to coexist and mutually support each other in the oral cavity, leading to a cross-kingdom synergistic pathogenic effect between these two microorganisms (25). Previous research has demonstrated that the *gtfB* and *gtfC* genes involved in the synthesis pathway of *S. mutans* through Gtfs play a role in promoting cross-kingdom interactions with *C. albicans*. Additionally, this study suggests that expression of *S. mutans* quorum sensing genes influences the formation of cross-kingdom biofilms (26). The phosphotransferase system and TCS are the primary pathways for regulating the production of EPS in *S. mutans. VicRK* is a specific type of TCS in *S. mutans*, where inhibition of the histidine kinase *VicK* and response regulatory transcription factors leads to negative regulation of Gtfs activity, resulting in reduced production of water-insoluble glucan and water-soluble glucan (27). LiaSR is another type of TCS in *S. mutans*, essential for adaptation and playing a crucial role in pH sensing and regulating cell membrane stress response. Deletion mutation in *liaR* results in continued production of LiaR by *liaS* (28). Our team’s prior research has shown a reduction in acid tolerance response in the liaS^-^ strain of *S. mutans* 593, with a similar trend observed in the *Streptococcus mutans* (NG-8) strain. However, there was no significant impact on acid sensitivity in *Streptococcus mutans* (UA 159). This is believed to be attributed to the varying roles of *liaS* across different strains. In the initial stages of this study, we screened *liaS* deleted mutation strains of UA 159 and *S. mutans* 593 (Supplementary Fig. 3).

Deletion of *liaS* mutation in *S. mutans* leads to abnormal cellular physiological regulation and altered expression of cariogenic function (22). This study also demonstrated that the role of *S. mutans*/*liaS* in cross-kingdom biofilms and its regulation of hyphal transformation in *C. albicans*. Gram staining results revealed that the cell chain-length of liaS^-^ was shorter than that of *S. mutans* 593, suggesting a potential impact on virulence function expression in *streptococcus* and microbial adhesion to hydrocarbons (29). The cell chain-length of liaS^-^ was properly restored and expressed after supplementation with the *liaS* gene. *S. mutans* demonstrates its survival advantage and cariogenic effect by balancing acid production and acid resistance. When the environmental pH falls below 5.5, the acid production efficiency of *S. mutans* significantly decreases and stabilizes at a pH of about 4.2. The survival rate of the *liaS* knock-out strain in an environment with a pH of 5.5 is significantly lower than that of the wild-type (30). Although *C. albicans* does not exhibit the same level of acid production efficiency as *S. mutans*, it is still capable of maintaining its acid production ability even in an environment with a pH=4.0 (31). The detection results from the biofilm model in vitro demonstrated that liaS^-^ significantly inhibited the pH drop in both single-and dual-species biofilm microenvironments. Furthermore, the *S. mutans* and *C. albicans* biofilm was found to promote glycolysis, organic acids, and EPS production (32). Additionally, the production of water-insoluble and water-soluble glucan in liaS^-^ was significantly inhibited due to decreased gene expression of *gtfB* and *gtfC* as confirmed by qRT-PCR analysis.

This study has demonstrated, for the first time, that *S. mutans*/*liaS* effectively enhances the virulence of *C. albicans* by promoting hyphal development and biofilm formation. When compared with *S. mutans* 593 and liaS^-^-comp, liaS^-^ formed smaller colonies and loose biofilm structures, with fewer bacteria adhered to the surface of fungi. Notably, *C. albicans* was predominantly found in the yeast form. The promotion effect of *liaS* on hyphal transformation was confirmed through GT formation and detection of Ergosterol content. The observed trend in *phr1* and *phr2* gene expression regulating pH adaptation in *C. albicans* was consistent with pH measurements. *S. mutans* induces hyphal transformation in *C. albicans* through the glucosyltransferase B (GtfB) regulatory pathway, where exogenous GtfB from S. mutans binds to fungal cells, promoting adhesion and sucrose decomposition into glucose and fructose. This process significantly upregulates genes and proteins related to carbohydrate metabolism in *C. albicans*, including sugar transport, aerobic respiration, pyruvate metabolism, and glyoxylate cycle (33).

The inhibition of glycolysis by *S. mutans* indirectly leads to a reduction in the production of GtfB, which promotes adhesion and metabolism of fructose and glucan on fungal surfaces to participate in regulating hyphal transformation (34). The deletion of *liaS* gene significantly downregulates the expression of the *gtfs* gene and reduces the adhesion of *S. mutans*. However, further research is needed to elucidate the specific regulatory mechanisms of the *liaS* gene on glycolysis and *gtfs* expression.

Oral candidiasis is closely associated with the virulence expression of *C. albicans*, and the inter-kingdom regulation of fungi by *S. mutans* depends on cell signaling transduction. Pre-treatment of the culture supernatant from *S. mutans* impacts the hyphal transformation, adhesion, and biofilm formation of *C. albicans* (35). The sigma factor *sigX* encoded by *S. mutans comX* is activated by *C. albicans* and regulates the reduction of Ras1-cAMP-Efg1 pathway expression to inhibit hyphal transition and fungal invasion (36). However, the promotional effect of GtfB secreted by *S. mutans* on EPS production and hyphal transformation appears to be more significant in cross-kingdom biofilms. Our study also revealed that *liaS* influences the expression of the *vicRK* two-component system and *comDE* quorum sensing system. *VicRK* regulates the Gtfs of *S. mutans* and can be activated by *C. albicans* to enhance its binding with GtfB (15, 37). Therefore, the production of GtfB is intricately based on interplay between different gene regulations in *S. mutans* and *C. albicans*, necessitating further investigations under varying conditions.

Additionally, the murine model of oropharyngeal candidiasis has shown that *S. mutans*/*liaS* significantly promotes fungi invasion. Deletion of *liaS* reduces virulence-enhancing effect of *S. mutans* on *C. albicans* by decreasing epithelial infection area, fungal burden, local epithelial damage, inflammatory infiltration, and hyphal invasion. Blocking the interaction between fungi and cells is an effective method for preventing and treating fungal infections caused by non-pathogenic or low-pathogenic expression of *C. albicans* in cross-kingdom biofilm formation (38). It is important to note that exogenous supplementation of cAMP restores the adhesion, hyphal transformation, and epithelial invasiveness of *C. albicans*. however, its role in *S. mutans* is unclear. In conclusion, targeted inhibition of *liaS* expression combined with cocktail therapy offers a promising treatment strategy for drug-resistant and microbial co-infections.

## Conclusion

Our study has shown that the deletion of the *liaS* gene effectively reduces EPS production in the cross-kingdom biofilm of *S. mutans* 593 and *C. albicans* and inhibits hyphal transition by downregulating the expression of the *gtfs* gene pathway and inhibit glycolysis efficiency in the biofilm. Additionally, *S. mutans* 593/*liaS* promotes epithelial invasion of *C. albicans* in murine oropharyngeal candidiasis, providing a potential therapeutic strategy for combating cross-kingdom infections caused by *S. mutans* and *C. albicans*.

## Materials and methods

### Microbial strains and growth conditions

All strains utilized in this study, including *C. albicans* SC 5314 and *S. mutans* 593, were sourced from the School of Stomatology at Fujian Medical University, China. The liaS^-^ strain denotes the *liaS* gene deletion mutant strain derived from *S. mutans* 593 (24). *C. albicans* was cultured overnight in Sabouraud dextrose broth (SDB, Solarbio Sci-ence&Technology Co., Ltd., China) at 35 °C (39). *S. mutans* was cultivated in Brain heart infusion (BHI, Oxoid, UK) medium at 37 °C with 5% CO_2_, while liaS^-^ was grown in BHI supplemented with 100 μg/ml spectinomycin (24).

The coculture of dual-strains is conducted in Tryptone yeast extract (TYE) medium, containing 25 g/L tryptone (Tryptone, Oxoid, UK), 15 g/L yeast extract (Yeast Extract, Oxoid, UK), and 1% (w/v) sucrose. The final concentrations of *C. albicans* and *S. mutans* in TYE are adjusted to 10^4^ CFU/ml and 10^6^ CFU/ml, respectively. For single-strain cultures and fresh medium, a 1:1 ratio is used; for dual-species cultures, half the volume of each microorganism is mixed with fresh medium. Cultures are maintained at 37 °C under 5% CO_2_ conditions (39).

### Construction of liaS^-^ complemented strain

Prepare the primers *liaS*-F and *liaS*-R (Supplementary Table) for amplifying the full-length *liaS* gene from the genomic DNA of *S. mutans* 593 (40). Subsequently, insert the gene fragment (SMU.486), encompassing the entire *liaS* open reading frame and promoter sequence, into the shuttle plasmid pIB169 to construct a linearized *liaS*/pIB169 plasmid. The Escherichia coli-streptococcal shuttle plasmid pIB169 was generously provided by Prof. Zhengwei Huang from Shanghai Jiao Tong University. Dilute the overnight culture of liaS-20-fold in BHI medium supplemented with 10% heat-inactivated horse serum (Sigma Aldrich, St. Louis, MO, USA). Following incubation at 37 °C for 4 h, add 1.0 μg/ml of competence-stimulating peptide (CSP, Shangon, Shanghai, China) and 10 μg of plasmid individually. Incubate the mixture at 37 °C for an additional 4 h and select the transformed strain of liaS^-^ with liaS/pIB169 on BHI agar medium containing 20 μg/ml chloramphenicol (41, 42). To validate the complementation of the liaS^-^ strain, chromosomal DNA extraction and qRT-PCR were performed to confirm *liaS* gene expression. The liaS^-^ complementary strain used in this study was monoclonally amplified in BHI with 20 μg/ml chloramphenicol.

### Biofilm formation and biomass assay

The 24 h biofilm is constructed in a 96-well plate and the CV staining method is employed to assess the formation of all biofilms. Subsequently, the supernatant culture medium is removed, and the biofilm is washed twice with PBS. Biofilms were fixed in methanol for 15 min and stained with 0.1% (w/v) CV for 10 min. After being washed with water and dried for 30 min, the biofilm sample is photographed using an optical microscope (ZEISS, Stemi 508). Added 33.33% (v/v) acetic acid to dissolve the biofilm. 100 μl of the mixed solution is taken out into a new well plate, and the biofilm is quantified on a microplate reader at OD575_nm_ (SpectraMax iD3). Biofilm samples are collected using sterile surgical blades and dissolved in PBS. The suspension containing cells is sonicated for 5 min to stir the aggregates. The single species and co-cultured biofilm gradients are diluted and evenly coated on SDA and BHI agar plates containing amphotericin B to quantify *C. albicans* and *S. mutans* in the biofilm, respectively. After incubation at 37 °C for 48 h, the biomass is evaluated. Each test is conducted in triplicate, independently three times, and the average log10 CFU is used to determine the fungal and bacterial biomass (43).

### Observation of biofilm structure

The biofilm structure, EPS production, and fungal morphology were examined using SEM. Circular sterile-glass slides with a radius of 0.6 cm were utilized as carriers for the biofilm. The biofilm was cultured in a 24-well plate, rinsed three times with sterile PBS, and then fixed with a 2.5% glutaraldehyde solution (J&K Scientific, Co., Ltd., China) (44). Subsequently, the biofilm was dehydrated in ethanol sequentially, dried, and sputter-coated with gold-palladium. SEM images were captured at magnifications of 2,000× and 4,000× in random fields of view using an EM8000 microscope (KYKY, China).

### Glycolysis and polysaccharide measurement

The MTT method was employed to determine the metabolic activity of biofilms (n=6/group). The construction method of biofilm is as described in CV staining and a final concentration of 0.5 mg/ml MTT solution was added and incubated for an additional 4 h at 37 °C. The formazan crystals were dissolved in 110 μl DMSO for 20 min. The absorbance of each sample was measured at OD490_nm_ (45).

According to the measurement of pH drop, modifications were carried out. To determine the effect of the *liaS* gene on glycolysis of *S. mutans*, single-strain and dual-species samples were collected in the logarithmic phase. Resuspended in 0.5 mM potassium phosphate buffer (containing 37.5 mM KCl and 1.25 mM MgCl_2_) and adjusted to 1% (w/v) by adding glucose. The Orion Dual Star pH/ISE Benchtop (Thermo Scientific; Waltham, Massachusetts, USA) was used to monitor the pH decrease at 5 min intervals for 120 min (46).

Collected single and dual-species biofilm samples (n=3) from 24 wells into 1.0 ml extraction solution. Sonicate the cells (in an ice bath, with a power of 200W, sonication for 3 seconds, an interval of 10s, and repeated 30 times), and centrifuge (8000 g, 4 °C, for 10 min) to extract the supernatant and place it on ice for detection. The pyruvate kinase (PK) assay kit (Solarbio, Beijing, China) was employed to detect the intracellular PK activity of *S. mutans*, and the changes in measurement values caused by NADH oxidation were measured at OD340_nm_ (35, 44). According to the manufacturer’s instructions, the PK activity is calculated as follows: PK activity (U/10^4^ cell) = [△A × Vtotal ÷ (ε × d) × 10^9^] ÷ (N × Vsample ÷ Vtotal) ÷ T = 2680 × △A ÷ N (Vtotal = the total volume of the reaction system, 2 × 10^−4^L; ε = the molar extinction coefficient of NADH, 6.22 × 10^3^L/mol/cm; D = the optical diameter of the 96-well plate, 0.6 cm; Vsample = the sample volume added, 1.0 ml; T = the reaction time, 2 min; N = the total number of bacteria, 10^4^.)

Water-soluble and water-insoluble glucans in the biofilms were analyzed using the anthrone-sulfuric method (39). Biofilm samples were collected by adding 1.0 ml ddH_2_O and centrifuging at 4 °C, 4000 rpm for 20 min, with this process repeated twice to obtain water-soluble polysaccharides. The biofilm was then resus-pended in 1.0 ml of 0.4 mol/L NaOH, followed by centrifugation at 4000 rpm for 20 min, with this step also repeated twice to collect the supernatant containing water-insoluble polysaccharides. Each supernatant sample (200 μl) was mixed with 600 μl of 0.2% (g/v) anthrone-sulfuric reagent separately and reacted in a 95 °C water bath for 10 min before being monitored by microplate reader at OD630_nm_ to calculate the glucans content in the biofilm according to the standard curve (Supplementary Figure 1).

### Hyphal transformation and Effect ergosterol synthesis

Researching the cross-kingdom interaction of *S. mutans*/*liaS* on hyphal transformation using the GT assay involves incubating the biofilm of *C. albicans* or cocultured biofilm at 37 °C for 24 h. After vortex thoroughly separates biofilm samples, the sample is dropped onto the Neubauer hemocytometer chamber and covered with a cover glass. Counting per 300 cells of *C. albicans* in the continuous field of view for each sample allows for calculation of hyphae morphologies cells at a magnification of 400× (47).

Ergosterol is a crucial component of the fungal cell membrane, characterized by its stable structure and strong specificity. The detection of ergosterol content can confirm differences in fungal growth and hyphae formation. Dual-species biofilm was collected, and the net weight was determined. Subsequently, 3.0 ml of 25% alcoholic potassium hydroxide solution was added to each sample, followed by mixing for 1 min using vertexing. The samples were then transferred to sterile borosilicate glass tubes and incubated in a water bath at 85°C. Afterward, 1.0 ml of ddH_2_O and a mixture of heptane were added to extract ergosterol from the samples. The extracted ergosterol was continuously measured at 230-300_nm_ using a spectrophotometer (UV-5200PC, Shanghai METASH instruments Co., Ltd., China) (48). The percentage of ergosterol + %24 (28) DHE was calculated using the formula [(A281.5/290) × F] / pellet weight for %24 (28) DHE = [(A230/518) × F] / pellet weight, %ergosterol = [%ergosterol + %24 (28) DHE]—%24 (28) DHE, where F represents the dilution factor in ethanol and E values are 290 and 518.

### RNA isolation and qRT-PCR

Collected dual-species biofilms were resuspended in 25 mg/ml lysozyme and incubated at 37 °C for 14 h to isolate RNA. TRIzol reagent (Invitrogen, Carlsbad, CA, USA) and chloroform were added, the supernatant was mixed with isopropanol after reaction and then centrifuged to obtain RNA (49). The RNA concentration and purity were measured using a NanoDrop 2000 spectrophotometer, and PrimeScript™ RT kit (TaKaRa Biotechnology, Japan) with gDNA Eraser was used to generate complementary DNA (cDNA).

Real-time quantitative polymerase chain reaction (qRT-PCR) was performed to assess the expression of *liaS*, as well as virulence genes such as the *S. mutans*-related *gtfB, gtfC*, and *gtfD*, and the *C. albicans*-related *als1, als3, hwp1, ece1, phr1*, and *phr2*. The TB Green™ Premix Ex Taq™ II kit (TaKaRa Biotechnology, Japan) was utilized for qRT-PCR following the manufacturer’s instructions. The PCR cycle was run in the StepOnePlus real-time PCR system (Applied Biosystem, CA, USA), with a program set at 30 s/95 °C and 40 cycles of 5 s/95 °C and 30 s/60 °C. Gene expression levels were normalized to internal reference genes *16S rRNA* and *18S rRNA* and relative fold changes were calculated using the 2^-ΔΔCT^ method. Primers used are listed in Supplementary Table 1 (Sangon Biotech,China).

### Murine oropharyngeal candidiasis model

Female BALB/c mice (16-20 g, 6 weeks old) were subcutaneously injected with cortisone acetate (3 mg per mouse in 200 μl PBS containing 0.5% Tween 80) and intraperitoneally injected with cyclophosphamide (100 mg/kg). On the second day after injection, the mice were anesthetized by intraperitoneal injection of 5% chloral hydrate (10 ml/kg). A swab pre-soaked in a mixture of 10^7^ CFU/ml *C. albicans* or 10^4^ CFU/ml *C. albicans* and 10^6^ CFU/ml *S. mutans* was placed on the tongue of a mouse for 30 min to observe the effect of *liaS* on oral candidiasis. Two days later, the weight of the mice was recorded, and then they were euthanized. The infected area of the tongue was observed using a stereoscope (Stemi 508, Zeiss, Germany). The tongue was longitudinally divided into two parts and homogenized in half before being cultured on SDA plates to quantify the count of *C. albicans*, while the other half was processed for histopathological analysis by staining tissue sections with hematoxylin/eosin (HE) and PAS to observe invasion and hyphal formation (9).

The animal experiments were performed in strict accordance with the guidelines of the Animal Welfare Act of the ethics committee of Fujian Medical University and meet the ethical requirements (license number IACUC FJMU 2024-0378). This experiment meets humane standards and minimizes the subject’s pain.

### Statistical analysis

All experiments were conducted in triplicate, individually. Significant effects of the variables were determined using one-way ANOVA, followed by SNK. Differences were considered statistically significant if *P* < *0*.*05*. Statistical analysis was performed using SPSS 16.0 software (SPSS Inc.; Chicago, IL, USA).

## Acknowledgments

We extend our sincere gratitude to Prof. Zhengwei Huang of Shanghai Jiao Tong University for generously supplying the *Escherichia coli*-shuttle plasmid pIB169.

## Funding

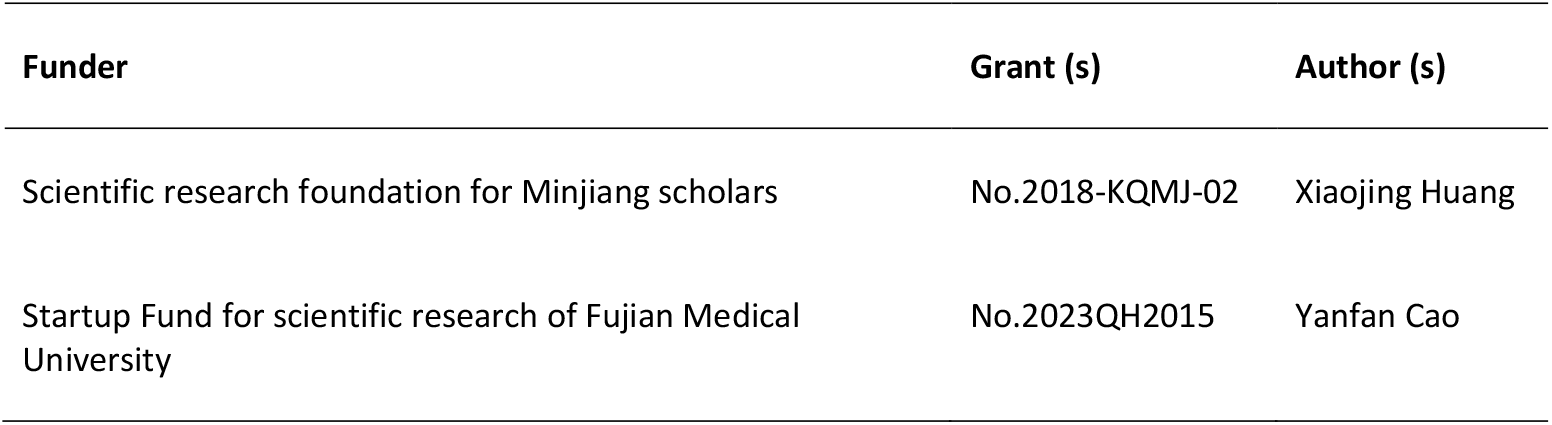

## Author contributions

Yanfan Cao, Conceptualized and designed the research, conducted experiments, and drafted the original manuscript for validation. Shan Huang and Jingyun Du, provided support with data analysis or theoretical framework. Xiuqing Wang, Guowu Gan and Shuai Chen, reviewed and made critical revisions to the manuscript. Xiaojing Huang, Secured funding, wrote, reviewed, and edited the manuscript, and contributed to conceptualization. All authors have read and approved the final version of the manuscript.

## Supplementary Material

**Table S1.**
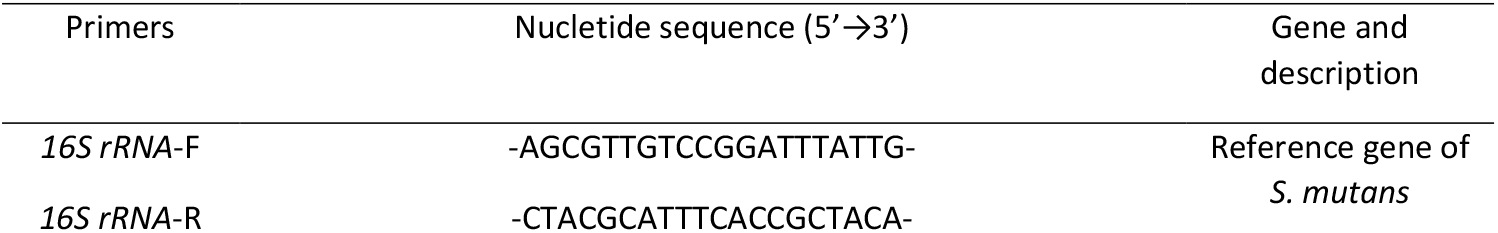
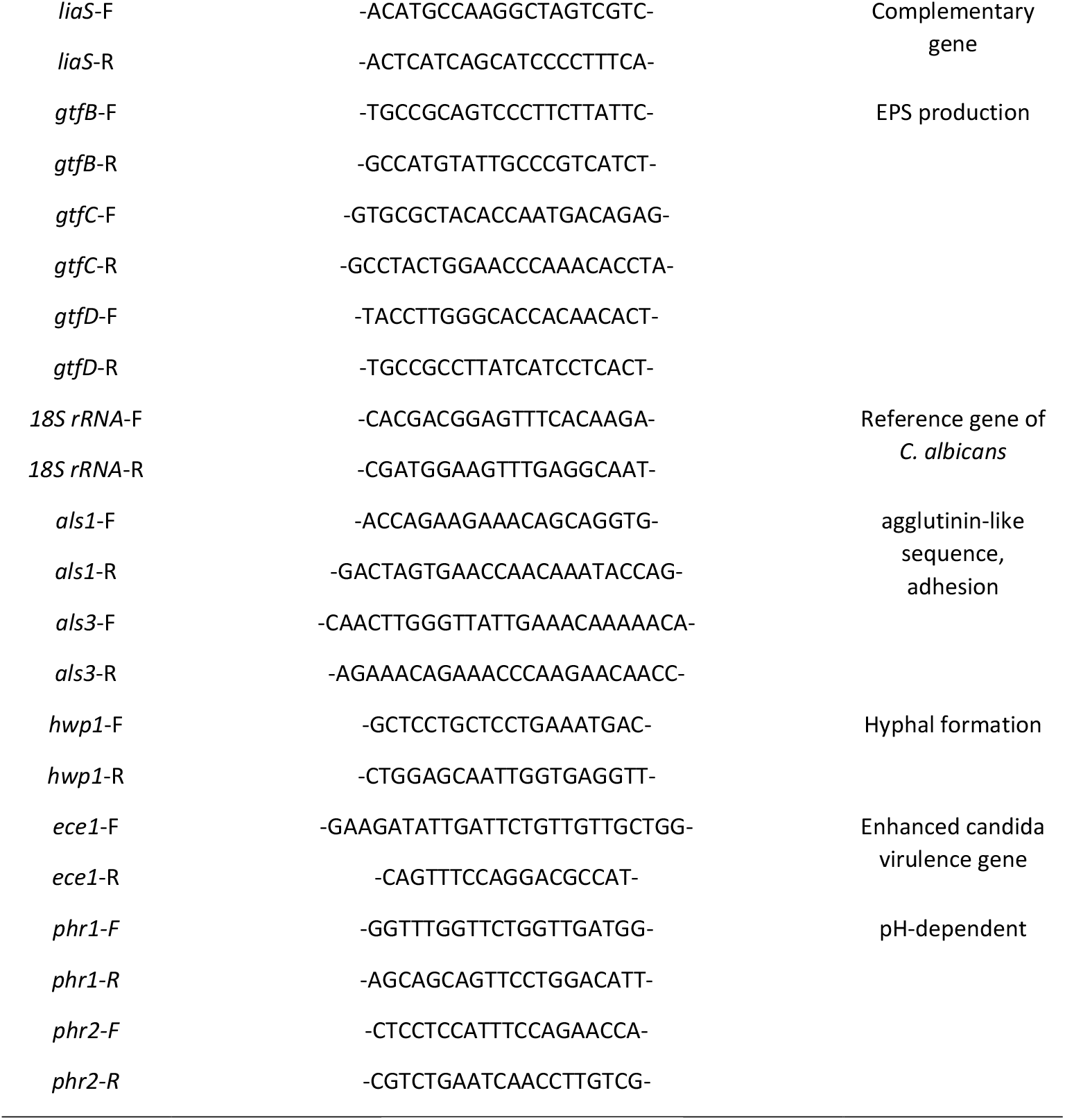

**Figure S1.**
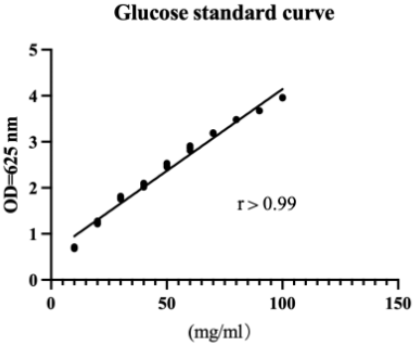
Determination of glucans standard curve at OD=625_nm_, which is used to measure the glucans content in biofilms.

**Figure S2.**
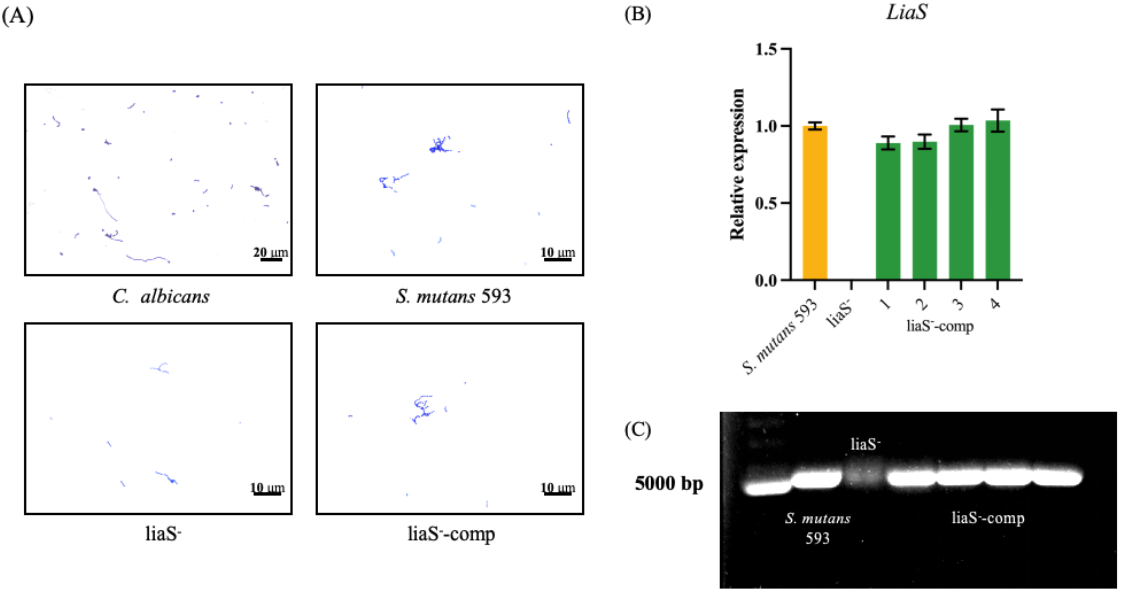
The morphologies of *C. albicans, S. mutans* 593, liaS^-^, and liaS^-^-comp was observed in Gram staining. The liaS^-^ strain exhibited a shorter chain-length compared to *S. mutans* 593 and liaS^-^-comp. (B)/(C) Gene and chromosomal DNA expression were analyzed using qRT-PCR and pulse-field gel electrophoresis for the screened strains. The liaS^-^ represents the *liaS* gene deleted mutation strain, and liaS^-^-comp represents the complementary strain. Experiments were conducted in triplicate and data are presented as mean ± standard deviations.

**Figure S3.**
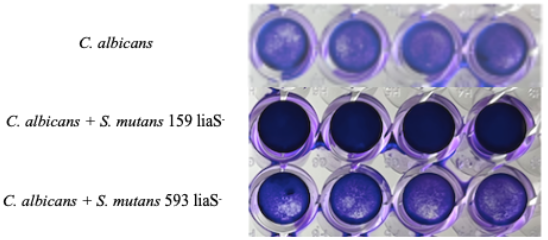
Crystal violet staining of biofilm in *C. albicans* and its co culture with *liaS* gene deleted mutant strains of *S. mutans* 593 and *S. mutans* 159 for 24 h cultivate.

